# Variations in terrestrial arthropod DNA metabarcoding methods recovers robust beta diversity but variable richness and site indicators based on exact sequence variants

**DOI:** 10.1101/693499

**Authors:** Teresita M. Porter, Dave M. Morris, Nathan Basiliko, Mehrdad Hajibabaei, Daniel Doucet, Susan Bowman, Erik J.S. Emilson, Caroline E. Emilson, Derek Chartrand, Kerrie Wainio-Keizer, Armand Séguin, Lisa Venier

## Abstract

Terrestrial arthropod fauna have been suggested as a key indicator of ecological integrity in forest systems. Because phenotypic identification is expert-limited, a shift towards DNA metabarcoding could improve scalability and democratize the use of forest floor arthropods for biomonitoring applications. The objective of this study was to establish the level of field sampling and DNA extraction replication needed for soil arthropod biodiversity assessments. Processing individually collected field samples recovered significantly higher richness (539-596 ESVs) than pooling the same number of field samples (126-154 ESVs), and we found no significant richness differences when using 1 or 3 pooled DNA extractions. Variations in the number of individual or composite samples or DNA extractions resulted in similar sample clustering based on community dissimilarities. Though our ability to identify taxa to species rank was limited, we were able to use arthropod COI metabarcodes from forest soil to assess richness, distinguish among sites, and recover site indicators based on unnamed exact sequence variants. Our results highlight the need to continue DNA barcoding of local taxa during COI metabarcoding studies to help build reference databases. All together, these sampling considerations support the use of soil arthropod COI metabarcoding as a scalable method for biomonitoring.

## Introduction

Soil arthropod fauna have been suggested as a key indicator of faunal community structure.^1–3^ These organisms are essential to ecological processes that include organic matter decomposition, nutrient cycling, and soil structural development (e.g. micropore formation that improves aeration porosity and water infiltration rates).^1,2^ Community shifts in soil arthropods in response to anthropogenic and natural disturbance have been documented in numerous studies.^3–7^

Typically, soil arthropods are sampled by trapping (e.g. pitfall traps) or they are extracted directly from soil (e.g. Tullgren funnels). Because of the large numbers of individuals that are sampled in even small studies, and because of the relative difficulty of identifying soil fauna, phenotypic identification is often expert- and time-limited. There are also significant issues of low recovery efficiency and bias in the recovery of soil fauna for phenotypic identification. A shift towards DNA metabarcoding could improve scalability and facilitate the use of soil arthropods for biomonitoring applications. DNA metabarcoding is currently the method of choice for highly scalable biodiversity studies.^8^ Although the use of COI metabarcoding to survey whole-community freshwater and Malaise trap arthropods is becoming fairly routine^9,10^, the use of COI metabarcoding to survey whole-community forest soil arthropods is still new.^11,12^

For DNA metabarcoding, bulk samples of soil are homogenized, DNA from all resident organisms are extracted, and a marker gene of interest is amplified using mixed template PCR. Marker genes are chosen according to the target organism, such as the cytochrome c oxidase subunit I (COI) mitochondrial DNA (mtDNA) marker that is the official animal barcode marker and has the largest number of reference sequences for taxonomic identification.^13,14^ This method produces ESVs (exact sequence variants) which are then compared to a reference sequence database. The reference sequence database is built through DNA barcoding of individual specimens identified using phenotypic characters.

The objective of this study was to establish the level of replication needed for field sampling and DNA extraction procedures for COI metabarcoding of terrestrial arthropods for biodiversity assessment. We assessed the influence of (1) increasing spatial sampling by including more individual or pooled samples (biological replicates), (2) performing mixed template PCRs on single or pooled triplicate DNA extractions (technical replicates), and (3) sampling from bryophyte, organic, and mineral layers (Fig S1) on observed richness, significance of sample clustering in beta diversity analyses, and the recovery of site indicators based on ESVs. We tested whether COI metabarcoding of forest floor arthropods could distinguish among two similar jack pine stands of differing origins.

## Results

### Sequencing Results

Raw sequence data was submitted to the NCBI SRA #XXX. A total of ∼ 41 million x 2 paired-end raw reads were sequenced for this project (∼ 110,000 reads per sample), of these ∼ 35 million (86%) raw reads were successfully paired, and of these ∼ 33 million (94%) paired reads were successfully primer-trimmed (Table S1). After primer trimming, the mode sequence length was ∼ 325 bp and ∼ 235 bp for the BE and F230 markers, respectively. A total of 67,626 denoised ESVs were detected where 19,562,246 primer-trimmed reads were mapped representing ∼ 47.5% of the original raw paired-end reads (Table S2). The phylum rank taxonomic distribution of the raw data is summarized in Fig S2. Only the 3,598 (4.8% of all ESVs) (BE 775; F230 2,823) ESVs that were assigned to Arthropoda were retained for further analysis below (Table S3). This corresponds to ∼ 2.7 million (6.5%) (BE 294,070; F230 2,398,638) of the original raw paired-end reads. Though the overall percentage of retained raw arthropod reads is low, it is consistent with previous work from bulk samples that are known to comprise a phylogenetically diverse mixture of taxa that are detected even when using primers originally developed to target arthropods.^15,16^ Since only a proportion of arthropod ESVs could be identified with confidence, we present our results at the ESV rank wherever possible (Fig S3). Rarefaction curves show that we saturated the sequencing of our arthropod COI PCR products (Fig S4).

### Effect of sampling method on richness

A total of 2,108 and 2,052 ESVs were detected from the Island Lake and Nimitz sites with some of the same ESVs detected across layers (Fig S5). ESV richness increases rapidly as more individually collected samples are added to the dataset (bioinformatically pooled samples), especially for the bryophyte and organic layers (Fig 1A). We also replicated this analysis using OTUs based on 97% sequence similarity with similar results as ESVs but with slightly lower richness values (not shown). Bryophyte and organic layer ESV richness also increases when more samples are manually pooled together, but at a lower rate than when individual samples are bioinformatically pooled (Fig 1B). The median richness detected from 15 individually collected, bioinformatically pooled, cores ranges from 539-596 ESVs and from 15 pooled cores ranges from 126-154 ESVs across sites. Since we used rarefaction to normalize the number of sequence reads included in these comparisons, we determined that the greater richness detected from individually processed cores compared with composited samples is due to the overall difference in the amount of soil sampled. For instance, a total of 33.75 g soil was extracted from 15 individually collected field samples (0.25 g x 3 layers x 3 DNA extraction replicates x 15 samples), compared with a total of 2.25 g soil from a composite of 15 pooled field samples (0.25 g x 3 layers x 3 DNA extraction replicates). We found that the ESV richness detected from a single individually collected field sample was not significantly different than processing a composite of 9-15 manually pooled samples. We did not test the effect of pooling DNA extractions across samples before PCR, or pooling PCR products across samples. There was also no significant difference in the ESV richness recovered when 1 or 3 DNA extractions were performed (Fig 1C and Fig S6).

**Fig 1.**
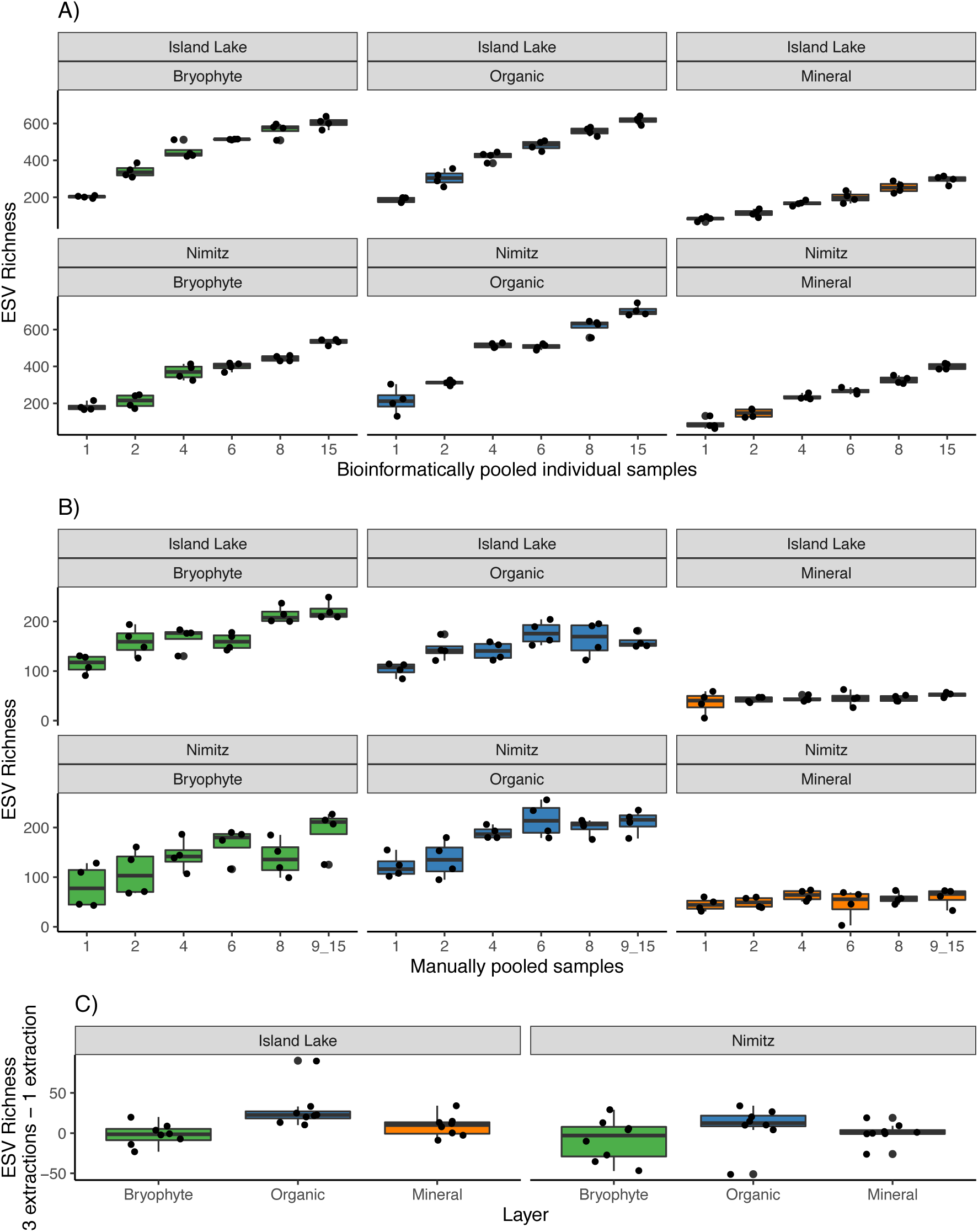
Arthropod ESV richness increases with increasing field sampling effort but varies little when more DNA extractions are performed. Richness is shown for A) bioinformatically pooled, individually collected field samples, B) manually pooled field samples, and C) the difference between samples processed with 3 pooled DNA extractions and 1 DNA extraction (positive values indicate greater richness from 3 pooled DNA extractions; negative values indicate greater richness from 1 DNA extraction). ‘915’ refers to the largest class of pooled cores that is 15 for all bioinformatically pooled samples but varies for manually pooled samples. At Island Lake the largest class is comprised of 15 pooled samples but at Nimitz, the largest class contains 15 pooled samples except for the bryophyte layer where 9-14 samples were pooled.

### Effect of sampling on beta diversity

Sample clustering across site and layer groups were similar across different methods (Fig 2). In our analysis of individually collected samples (Fig 2A), site and layer groups are clearly distinguished (NMDS: stress = 0.07, linear fit R^2^ = 0.96). We did not detect any significant beta dispersion (AVOVA: sites p = 0.67, layers p = 0.18) or interactions between site and layer groups (PERMANOVA: p = 0.06). In our analysis including samples derived from manually pooling increasing numbers of cores (Fig 2B) (NMDS: stress = 0.11, linear fit R^2^ = 0.99), we found significant beta dispersion among soil layer groups (ANOVA: experiment p-value = 0.86, sites p-value = 0.19, layers p-value = 0.01) but we proceeded with PERMANOVA because we had a statistically balanced design.^17^ We did not detect any significant interactions between site, layer, or number of manually pooled samples (PERMANOVA, p-value > 0.05). In our analysis including samples processed with 1 or 3 DNA extractions (Fig 2C) (NMDS: stress = 0.12, linear fit R^2^ = 0.93), we found significant beta dispersion only among layer groups (ANOVA: experiment p-value = 0.89, sites p-value = 0.74, layers p-value = 0.09), and we did find a significant interaction among site, layer, and experiment groups (p-value = 0.03). PERMANOVA results show that sites, layers, and method differences explain only a small but significant amount of the variation among samples, suggesting that other environmental factors likely drive community dissimilarities in these samples (Table 1).

**Table 1.** Amount of variation of beta diversity explained by groups varies according to sampling method.

| Experiment | PERMANOVA Formula | Group | F.model | R<sup>2</sup> | P-value |
| --- | --- | --- | --- | --- | --- |
| Individual field samples | site*layer | site | 18.9193 | 0.08243 | 0.038* |
|  |  | layer | 1.4108 | 0.01229 | 0.059 |
|  |  | site:layer | 1.3864 | 0.01208 | 0.055 |
| Manually pooled field samples | site*layer*expt | site | 1.1129 | 0.00751 | 0.020* |
|  |  | layer | 5.2626 | 0.07103 | 0.001* |
|  |  | expt | 1.3034 | 0.04398 | 0.056 |
|  |  | site:layer | 1.0515 | 0.01419 | 0.325 |
|  |  | site:expt | 0.7989 | 0.02696 | 0.908 |
|  |  | layer:expt | 0.9817 | 0.06625 | 0.512 |
|  |  | site:layer:expt | 0.7113 | 0.04800 | 0.997 |
| One versus three pooled DNA extractions | site*layer*expt | site | 0.5036 | 0.00498 | 0.001* |
|  |  | layer | 1.5000 | 0.02969 | 0.047* |
|  |  | expt | 9.3499 | 0.09254 | 0.001* |
|  |  | site:layer | 1.0568 | 0.02092 | 0.371 |
|  |  | site:expt | 0.4628 | 0.00458 | 0.998 |
|  |  | layer:expt | 1.1707 | 0.02317 | 0.206 |
|  |  | site:layer:expt | 1.6319 | 0.03230 | 0.030* |

**Fig 2.**
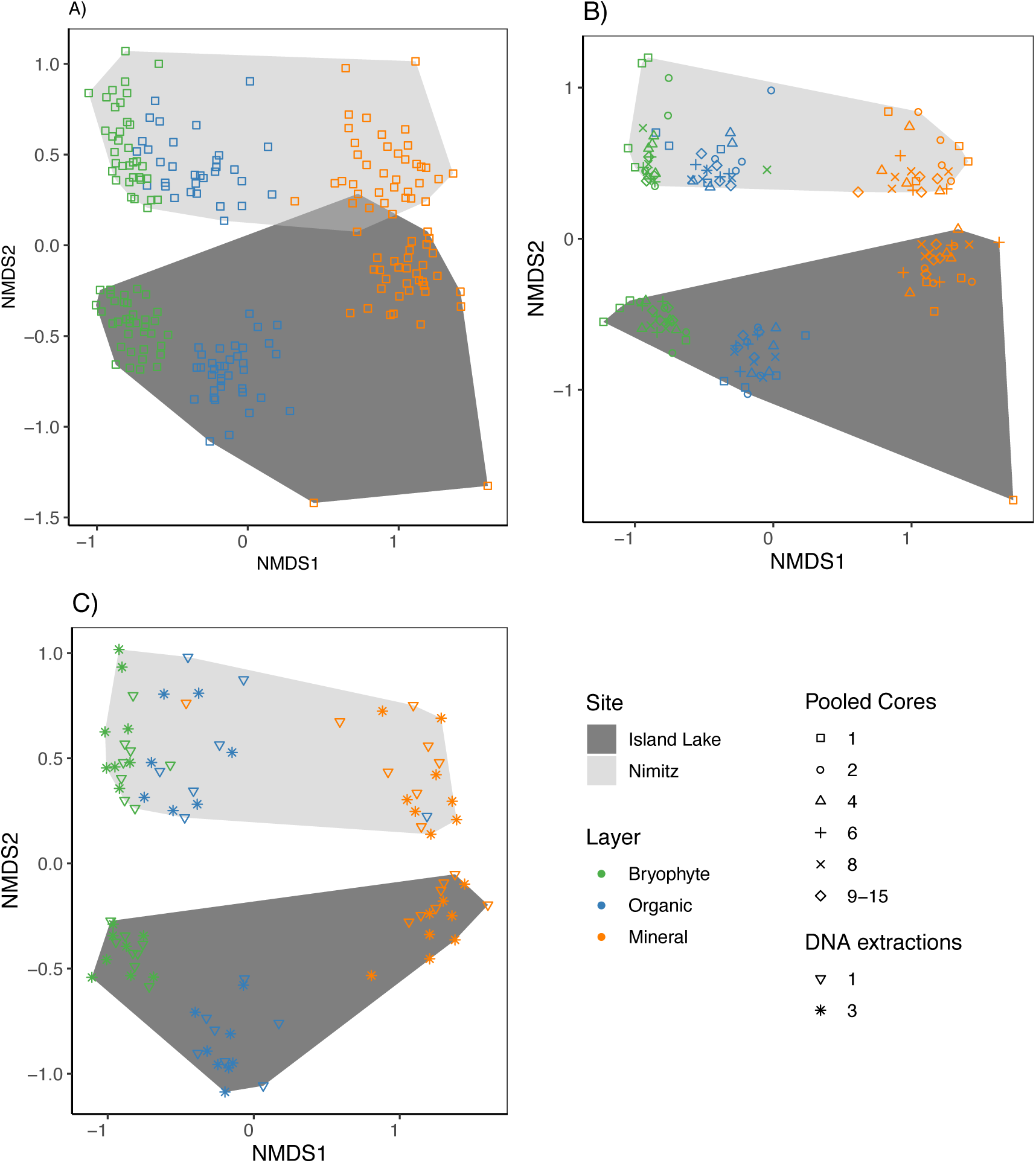
Clustering of samples across sites and soil layers are robust to intensity of field sampling and number of DNA extraction replicates. Clustered groups are shown based on A), the collection of individually processed samples, B) manual pooling of 1-15 samples and C) single samples processed with 1 or 3 pooled DNA extractions.

The group ‘site’ refers to Island Lake or Nimitiz field sites; ‘layer’ refers to bryophyte, organic, or mineral layers; ‘expt’ refers to variation in the sampling methods such number or pooled cores or the number of pooled DNA extractions. The asterisk (*) indicates significant results.

### Assessing the stability of site indicator analyses

Higher level, the most inclusive, taxonomic composition of site indicator ESVs is similar across sties (Fig 3). The fine level taxonomic composition of site indicator ESVs is not always resolved to the species rank because of our inability to make high confidence taxonomic assignments. For improved readability, we plotted heat trees summarized to the species rank although site indicator analysis was conducted using ESVs. Where the same indicators appear to be detected from both sites, this is often due to our inability to confidently identify the ESVs, but the variation at the ESV level of resolution can distinguish among these sites (Fig S7). Soil arthropods from both sites were comprised of mainly Arachnida (Scorpiones, Araneae, Sarcoptiformes), Insecta (Trichopitera, Hemiptera, Hymenoptera, Lepidoptera, Coleoptera, Diptera), Collembola (Entomobryomorpha), Malacostraca (Decapoda), and Diplopoda (Polydesmida). Many site indicator taxa are detected infrequently among samples. For the Island Lake site, site indicator ESVs from unknown Trombidiformes (plant parasitic mites), *Oppia nitens* (polyphagous fungiverous mite), *Eniochthonius crosbyi* (mite), unknown Plecoptera (stoneflies), Odonata (carnivorous dragonflies/damselflies), unknown Orthoptera (herbivorous grasshoppers/locus/crickets), Entomobryidae (omnivorous slender springtails), *Folsomia nivalis* (elongate-bodied springtail), and unknown Poduromorpha (springtails) were found in more than half the samples. For the Nimitz site, indicator ESVs from unknown Siphonaptera (parasitic fleas), unknown Phasmatodea (herbivorous stick insects), and *Isotoma riparia* (springtail) were found in more than half the samples. We also illustrate how the taxonomic composition of site indicator ESVs varies slightly according to the number of manually pooled field samples, but in no consistent way (Fig S8); and varies minimally according to the number of DNA extractions used (Fig S9). However, we did find that taxonomic diversity of site indicator ESVs across soil layers was quite variable, with the majority of indicator ESVs recovered from the bryophyte and organic layers (Fig S10).

**Fig 3.**
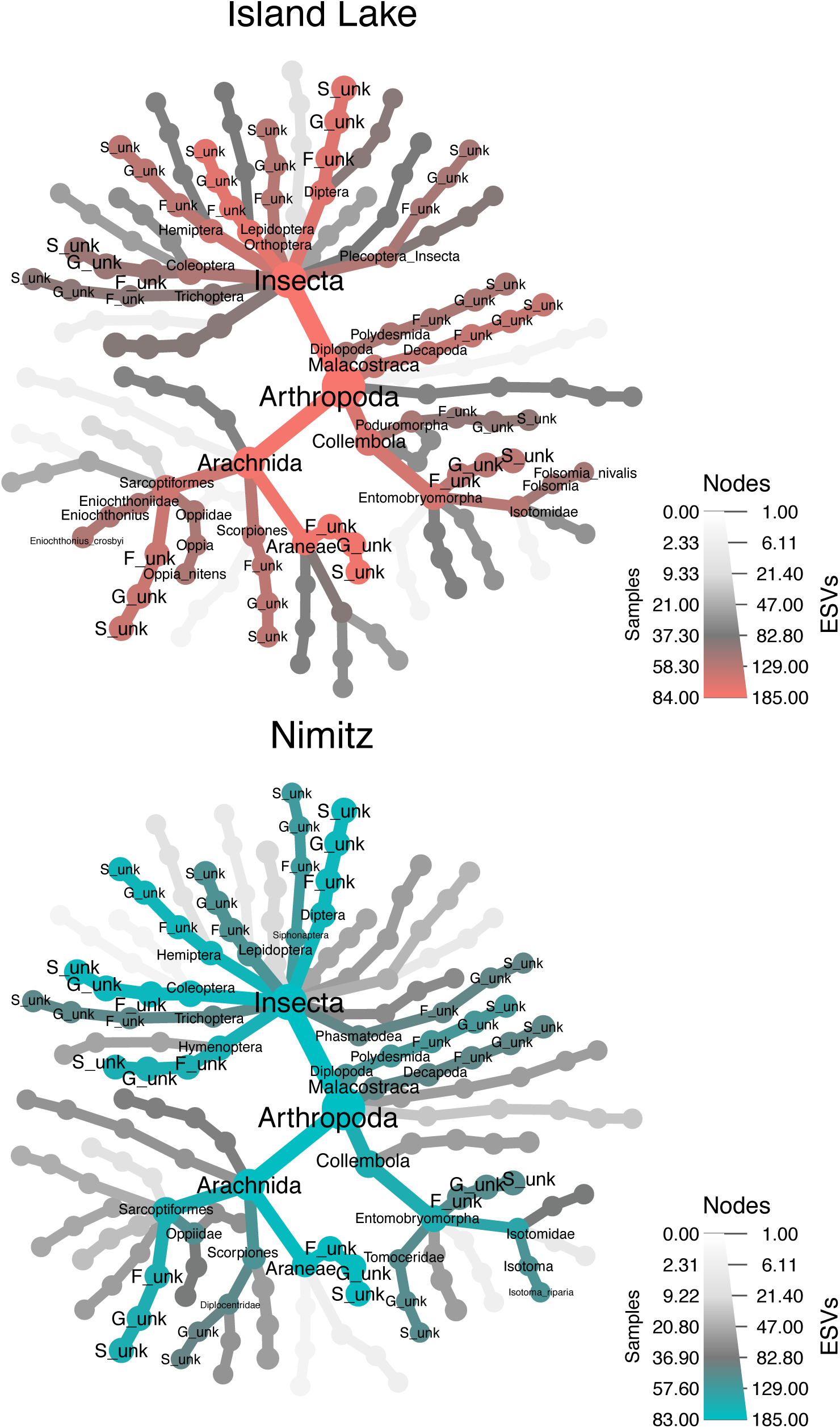
Taxonomic distribution of site indicator ESVs for each site. Heat trees comprised of all the site indicator ESVs, pooled across all sampling methods, are shown for each site. In each tree, color indicates the frequency of taxa detected across samples; text and node size indicate the number of indicator ESVs that are summarized in each taxon. To improve readability, labels have been added only to nodes present in at least half the samples. Taxa that could not be confidently identified are indicated as follows: F_unk = family unknown, G_unk = genus unknown, S_unk = species unknown.

## Discussion

Our results highlight that the inclusion of replicate soil samples is critical to detect maximum richness of arthropods that have patchy spatial distributions. When we conducted richness calculations based on either ESVs or OTUs, results based on OTUs showed slightly lower richness but the trends were similar, i.e., increasing numbers of pooled samples resulted in a greater number of sequence clusters detected. For ease of bioinformatic reproducibility and comparability across studies, the use of exact sequence variants has been encouraged by others and was the method adopted in the current study.^18^ Both ESVs and OTUs have been shown to perform similarly in biodiversity analyses when calculating richness and beta diversity.^19^ When analyses at a certain taxonomic rank are needed, both ESVs and OTUs can be taxonomically assigned. We also found that the comparison of beta diversity across sites is robust to variations in field sampling methods. Changes in the number of pooled DNA extractions from the same sample also produced similar results with respect to richness and beta diversity. If resources are limited, a single DNA extraction per sample would be sufficient to process well-homogenized soil samples. Our results complement a previous study conducted across grassland, forest, and cropland sites where differences in sampling methods (conventional morphology, DNA metabarcodoing of bulk soil and extracted arthropods) resulted in differences in the detection of individual taxa, but yielded similar site level diversity and composition.^11^ Our results are also consistent with a previous simulation study that used an earthworm dataset to show how multiple samples from the same location sometimes recovered slightly different communities but multiple DNA extractions from the same sample accurately detected the target taxa.^20^ With limited resources available, it would be more effective to put more effort into replicating sampling at the field site level, than it is to spend the time manually pooling field samples or performing replicate DNA extractions. We did not, however, test the effect of collecting many replicate field samples and sequencing them individually (done here) versus pooling increasing numbers of single sample DNA extracts prior to mixed-template PCR, or pooling increasing numbers of PCR products prior to sequencing. These types of experiments would provide guidance on the design of future soil arthropod surveys to help keep the cost of molecular biology supplies and sequencing per sample to a minimum.

Richness, beta diversity, and indicator taxon analyses show differences across soil layers. The higher arthropod richness we observed in the bryophyte and organic layers is consistent with results from another Island Lake study that used phenotypic classification of Collembola (springtails) and Oribatida (mites).^3^ In the Rousseau et al., 2018 study, they showed higher density, biomass, and diversity of springtails and mites in moss and organic soils compared mineral soil. This has important implications with respect to sampling strategy and suggests that separating samples by soil horizon is a critical consideration for generating comparable samples between sites. In addition, this horizon separation supports the hypothesis that the moss layer is a critical resource for arthropods and its recovery after disturbance is likely necessary for a return of mature forest soil faunal communities.^3^ In our study sites, minimal diversity would be missed if the mineral layer was not sampled, but future work should test this across a broader range of forest soils. These sampling considerations support the use of soil arthropod CO1 metabarcoding as a scalable method for biomonitoring.

We know that current COI reference databases such as BOLD and GenBank are not complete, fortunately database representation has been shown to be improving year after year.^21,22^ This limitation does have implications for studies working to benchmark DNA metabarcoding protocols against previous work based on commonly used bioindicator species. False negatives, taxa missed by DNA metabarcoding, can occur when local species have not yet been DNA barcoded and are missing from the reference sequence databases.^11,23^ For example, when we compared the species list from the Rousseau et al.2018 study also conducted at Island lake with the taxa present in the COI classifier v3, we found that 70% of their fully identified springtail and mite species (36% of genera) were missing from the reference database. This further highlights the importance of supplementing CO1 metabarcoding studies with local DNA barcoding to improve taxonomic assignment rates.^24^

Site comparisons and the detection of site indicators using soil arthropod metabarcodes, however, can still be conducted whether or not the sequence clusters have been taxonomically assigned. In this study, we showed beta diversity comparisons using ordination and PERMANOVA that successfully distinguished samples among sites without using any of our taxonomic annotation data except for some upfront filtering of the dataset for arthropoda sequences. We also showed how site indicators based on exact sequence variants were successfully recovered even though taxonomic assignments to more inclusive levels of resolution appeared similar across sites. Our results are consistent with a previous study that showed how COI metabarcoding may actually recover a greater taxonomic diversity of site indicators in addition to the usual expected bioindicators.^15^ Future studies should attempt to pair soil arthropod metabarcoding with local DNA barcoding to improve taxonomic assignment rates. Despite this, the use of soil arthropod metarbarcodes as site bioindicators was successful and samples from two similar jack pine stands with different origins were distinguished from each other based on beta diversity and the presence of site indicators.

## Methods

### Study Area and Field Sample Collection

Moss and soil samples were collected from 2 boreal forest stands in north-central Ontario that differ in origin. The first site is a 51-year old jack pine (*Pinus banksiana*) stand that was previously clearcut and located at the Island Lake Biomass Harvest Research and Demonstration area approximately 20 km from Chapleau, Ontario, Canada (47° 42’ N, 83° 36’ W).^25^ The second site was a 92-year old jack pine stand of wildfire origin (47° 38’ N, 83° 15’W). Mean annual temperature and precipitation for the area is 1.7°C and 797 mm (532 mm of rainfall and 277 cm of snowfall), respectively (Environment Canada 2013). These two jack pine-dominated stands (>90% jack pine, based on live tree basal area) were established on glaciofluvial, coarse-textured, glacial outwash deposits characterized by sandy (medium sand) parent material overtopped with a variable depth loess (windblown) cap of finer textured soil (silty fine sand to silt loam).^26^ They both have a moderately dry soil moisture regime with rapid drainage. Forest floor depth (i.e. LFH – Litter, Fermented, Humic) was approximately 9-10 cm.

At each site we chose a 2 m × 2 m area of continuous moss cover (Fig S1). Starting in the northwest corner, we used a bread knife to cut a 5 cm × 10 cm × full depth volume of moss and placed it in a labeled zip top bag. We sterilized the knife and spoon between samples by cleaning with 70% ethanol. We then used a spoon to sample a 5 cm × 10 cm x full depth volume of the organic horizon (LFH) and placed it in a labeled zip top bag. The top 10 cm of the mineral horizon was sampled by hammering a 5 cm diameter × 10 cm long piece of polyvinyl chloride (PVC) pipe into the mineral horizon, extracting it, and placing it in a labeled zip-top bag. We repeated this procedure in a 6 × 6 grid (36 samples in total) with 20 cm spacing between each sample. In total we had 36 samples, with a subsample from each layer, at each site, for a total of 216 samples. Samples were immediately placed in a cooler with ice packs and were frozen at -20C within several hours of collection.

### Sample preparation

The wet weight was obtained for each sample. Bryophyte and organic samples were separately homogenized using a knife mill and mineral samples were homogenized by forcing them through a 0.2 mm sieve (Fig S9). The knife mill and sieve were both rinsed with water and then cleaned with 70% ethanol between samples. Samples from different soil layers were always kept separate. To thoroughly sample the soil arthropod community, samples were processed by subsampling 0.25 g of soil 3 times, extracting DNA from each replicate, and then pooling the DNA prior to PCR (1C3E method, 1 core, 3 DNA extractions). To assess the influence of using samples drawn from increased spatial sampling of soil, we subsampled 1 g of soil from each homogenized bryophyte and organic sample and 5 g of mineral soil to create composites drawn from each of 2, 4, 6, 8, and 15 samples (keeping layers separate). Each composite sample was put into a zip-top bag and shaken by hand to mix. This pooling was replicated 4 times with different samples represented in each pool (XC3E method, 2-15 pooled cores, each with 3 DNA extractions). For the Nimitz site, there was not always enough bryophyte sample so the largest pool had anywhere from 9 to 14 samples included. We used 0.25 g from each pooled soil sample for triplicate DNA extractions that were pooled prior to PCR. To assess the value of pooling multiple DNA extractions per sample, 0.25 g from each of 8 un-pooled soil samples x 3 soil layers were extracted one time only prior to PCR (1C1E method, 1 core, 1 DNA extraction).

### Molecular biology methods

DNA extraction was carried out using the DNeasy PowerSoil DNA Isolation Kit (Qiagen Cat# 12888-100) modified with the Braid et al. (2003) protocol that uses a chemical flocculant to help remove soil-derived PCR inhibitors.^27^ We extracted DNA from 0.25 g of soil per sample following the manufacturer’s protocol except that 200 μl of 100 mM aluminum ammonium sulfate dodecahydrate was added to the tube with 60 μl of solution C1 followed by a 10 minute incubation at 70°C to help lyse difficult samples.

Mixed template PCR and Illumina library preparation was carried out at the Canadian Forest Service’s Laurentian Forestry Centre. DNA was quantified using the Qubit dsDNA HS Assay Kit (Life Technologies, Burlington, ON, Canada). DNA concentrations were standardized to 5 ng/μl for all samples and each sample was amplified in triplicates to ensure reproducibility.^28,29^ Invertebrate communities were targeted using two sets of primers targeting the COI gene (5’- >3’): the F230R_modN marker with the forward primer LCO1490 GGTCAACAAATCATAAAGATATTGG and the reverse primer 230R_modN CTTATRTTRTTTATNCGNGGRAANGC adapted from the Gibson et al. 2014 230R primer to include N’s instead of inosines^30,31^; and the BE marker with the B forward primer CCIGAYATRGCITTYCCICG and the E reverse primer GTRATIGCICCIGCIARIAC.^9^ These primers were combined with the required Illumina adaptors at the 5′ end of the primer sequences, TCGTCGGCAGCGTCAGATGTGTATAAGAGACAG for the forward primer and GTCTCGTGGGCTCGGAGATGTGTATAAGAGACAG for the reverse primer. PCR reactions were set up by creating a master mix of 37.5 μl of HotStarTaq Plus Master Mix (QIAGEN Inc., Germantown, MD, USA), 1.5 μl of each 10 μM primer, 27 μl of UltraPure™ DNase/RNase-Free Distilled Water (GIBCO, Life Technologies) and 7.5 μl of gDNA at 5 ng/μl. The final volume of 75 μL was then distributed in three 96-well plates placed in separate thermocyclers. Thermal cycling conditions were as follows: initial denaturation at 95 °C for 5 min; 40 cycles at 94 °C for 30 s, 50 °C for 30 s, 72 °C for 1 min; and a final elongation at 72 °C for 10 min. Triplicates PCR products were pooled and visualized on GelRed-stained 1% agarose gels using the Chemigenius Bioimaging System (Syngene, Cambridge, UK). PCR products were purified using 81 μl of magnetic bead solution (Agencourt AMPure XP, Beckman Coulter Life Science, Indianapolis, IN, USA) according to Illumina’s protocol.^32^ Indexes were added to each sample by amplifying 5 μl of the purified PCR product with 25 μl of KAPA HIFI HotStart Ready Mix, 5 μl of each Nextera XT Index Primer (Illumina Inc., San Diego, CA, USA) and 10 μl of UltraPure™ DNase/RNase-Free Distilled Water for a total volume of 50 μl. Thermal cycling conditions were as follows: 3 min at 98 °C, 8 cycles of 30 sec at 98 °C, 30 sec at 55 °C, 30 sec at 72 °C, and a final elongation step of 5 min at 72 °C. Indexed amplicons were purified with the magnetic beads as previously described, quantified using a Qubit dsDNA BR Assay Kit (Life Technologies) and combined at equimolar concentration. Paired-end sequencing (2 × 250 bp) of the pools was carried out on an Illumina MiSeq at the National Research Council Canada, Saskatoon.

### Bioinformatic methods

Reads were processed using the SCVUC v2.0 bioinformatic pipeline available from GitHub at xxx. SCVUC is an acronym that stands for the major programs/algorithms used in the pipeline: “S” SEQPREP, “C” CUTADAPT, “V” VSEARCH, “U” USEARCH-unoise, “C” COI Classifier. At certain points commands were run in parallel using GNU parallel.^33^ First, the compressed fastq raw reads were paired with SEQPREP using the default parameters except that we required a minimum Phred score of 20 in the overlap region and a minimum overlap of at least 25 bp. Primers were trimmed with CUTADAPT v1.10 with the default settings except that we required a minimum length (after trimming) of at least 150 bp, a minimum Phred score of 20 at the ends, allowing a maximum of 3 Ns. CUTADAPT was also used to convert the compressed fastq files to compressed FASTA files. We added the sample name to the FASTA headers and concatenated all the sequences into a single file to permit the generation of global ESVs below. Sequences were dereplicated with VSEARCH v2.5.0 with the --derep_fulllength command, sequences comprised of identical substrings are retained as unique sequences, and the number of reads in each cluster were tracked with the --sizein --sizeout commands. Unique sequences were denoised and a set of ESVs were generated with USEARCH v10.0.240 with the unoise3 algorithm. With this method, predicted sequence errors are corrected, putative PhiX contamination is removed, putative chimeric sequences are removed, and rare ESVs are removed. We defined rare ESVs as clusters containing only one or two reads (singletons and doubletons) because it has been shown that rare clusters tend to be predominantly comprised of reads with sequence errors.^34,35^ In total, 41% of primer-trimmed reads belonging to rare ESVs were removed after denoising. To compensate for a known bug in this version of the program, we changed the ‘Zotu’ prefix in the FASTA file headers to ‘Otu’. At each major step of bioinformatic processing above, statistics including read/cluster number and read length (min, max, mean, median, mode) were calculated. Due to the limitations of the USEARCH 32-bit program, we used VSEARCH to construct the ESV x sample table that tracks read numbers in the ESVs. This was done by mapping good quality primer-trimmed reads to the denoised ESVs with 100% sequence similarity. At this step, shorter sequence substrings may be mapped to longer ESVs. The denoised ESVs were taxonomically assigned with the COI Classifier v3 available from https://github.com/terrimporter/CO1Classifier. Read number and samples were mapped to the taxonomic assignment table. We identified high confidence taxonomic assignments using the recommended minimum bootstrap cutoff values for 200bp fragments (species >= 0.70, genus >= 0.30, family >= 0.20). Assuming that our taxa are in the reference database, then taxonomic assignments should be at least 99% correct (95% correct for species).

To assess the stability of results at varying levels of resolution, we compared results based on ESVs and OTUs (operational taxonomic units). Denoised ESVs from above were fed into the ‘--cluster_smallmem’ command in VSEARCH and OTU clusters based on 97% sequence similarity were generated. These results were then processed as described above for ESVs, except that ‘-- id 0.97’ was used to map reads to the sample x OTU table.

### Data analysis

The BE and F230 taxonomy tables were prepared at the command line, with Perl, and analyzed in R v3.4.3 with scripts available from GitHub at xxx.^36^ The ‘vegan’ v2.4-6 package in R was used to plot rarefaction curves using the ‘rarecurve’ function.^37^ Rarefaction to the 15^th^ percentile was performed in vegan with the ‘rrarefy’ function. This was done to minimize library size bias in diversity comparisons.^38^ We then transformed read abundances to presence-absence data. We did this because PCR primer bias may distort template to PCR product ratios making read number unsuitable for inferring quantitative differences in biomass, density, or community composition.^39–41^

We assessed the effect of increasing spatial sampling by including more individual samples. We simulated sampling increasing numbers of individual cores by randomly bioinformatically pooling data from 1 – 15 individually collected cores and replicated this sampling 4 times. We calculated richness for each level of sampling effort using the ‘specnumber’ function in vegan. We calculated venn diagrams using the ‘vennCounts’ function in the limma Bioconductor package and plotted this using the ggforce package to draw circles.^42,43^ We assessed the effect of sampling effort on beta diversity using non-metric multi-dimensional scaling (NMDS) ordination. The NMDS plot was created with the ‘metaMDS’ function in vegan with 2 dimensions using Bray-Curtis dissimilarity with binary data (Sorensen dissimilarity). The number of dimensions was chosen by calculating a scree plot using the ‘dimcheckMDS’ function in the goeveg package (not shown).^44^ A Shephard diagram and goodness of fit calculations were created using the ‘stressplot’ and ‘goodness’ functions in vegan. Beta dispersion was assessed using the ‘betadisper’ function in vegan. We tested for significant interacting factors with permutational multivariate analysis of variance (PERMANOVA) using the ‘adonis’ function in vegan with the strata option so randomizations occur within sites.

We also assessed the effect of increasing spatial sampling by manually pooling increasing numbers of cores. For a balanced design, we randomly subsampled cores from the 1C3E method down to 4 replicates to match the number of replicates available for the XC3E method. We calculated richness for each level of sampling effort as described above. Beta diversity was assessed as described above (n=145), a single outlier was identified, removed, then the analysis was re-run.

We assessed the effect of performing mixed template PCRs on single or pooled triplicate DNA extractions. For a balanced design, we subsampled cores from the 1C3E experiment to match the same 8 grid coordinates as used in the 1C1E experiment. We calculated richness and beta diversity as described above. We calculated the difference in richness from the same samples processed using one or three DNA extractions, checked for normality using Shapiro-Wilk’s test for normality and tested for significant differences in richness across cores using pairwise Wilcox tests and adjusting p-values for multiple comparisons using the Benjamini & Hochberg (1995) method.^45,46^ Beta diversity was assessed as described above (n=95), two outliers were identified, removed, then the analysis was re-run. Since we detected a significant interaction between layer and experiment groups in the PERMANOVA, we tested for significant pairwise interactions using the pairwise.adonis function with Bonferroni p-value correction.^47^

Site indicator ESVS were determined using the ‘indicspecies’ v1.7.6 package in R using the multipatt command.^48^ Briefly, the indicator species concept describes the species associated with a certain site or condition based on their fidelity to those conditions and absence from others. This concept can be extended to the ESV or OTU rank when current reference databases do not allow us to identify all sequences to the species rank. To create a balanced design, a subsample from the same 4 grid coordinates from the 1C1E and 1C3E methods was used to compare with the 4 replicates available from the XC3E method. For each sampling method, site indicators were retained if they had a p-value <=0.05. To check whether the presence of recovered site indicators were similar across experiments, we calculated Pearson correlations with the corr.test function and adjusted for multiple testing using the “holm” method with the corr.p function in the ‘psych’ package.^49^ Significant correlations were visualized using the ‘corrplot’ v0.84 package in R.^50^ To illustrate the taxonomic distribution of site indicators and their prevalence across samples, the ‘metacoder’ v0.3.0 package in R was used to create heat trees.^51^An ESV x sample matrices enumerating the reads recovered for site indicator ESVs using each method were formatted in R to resemble QIIME output. From this, a sample matrix was constructed in R. To improve clarity, we reduced the number of edges in the heat trees by summarizing ESV taxonomic assignments to the species rank (instead of the ESV rank). The taxonomic information was parsed using the parse_tax_data command from the ‘tax’ v0.3.1 package in R.^52^ Taxon abundance at all ranks was calculated with the calc_taxon_abund command. Taxon occurrence per sample group was calculated with the calc_n_samples command.

## Data Accessibility

Raw reads are available from the NCBI Short Read Archive (SRA) (xxx). A FASTA file of final ESVs and the taxonomy table are available in the Supplementary Material. The SCVUC v2.0 bioinformatic pipeline is available from GitHub at xxx. The scripts used to produce figures are also available from GitHub at xxx.

## Acknowledgements

T.P. and L.V. would like to acknowledge funding from the Canadian government through the Genomics Research and Development Initiative (GRDI), EcoBiomics project. We would like to acknowledge Rob Fleming and Paul Hazlett at the Canadian Forest Service’s Great Lakes Forestry Centre for the development of the study area. We also appreciate the contribution by Christine Martineau and Marie-Josée Morency at the Canadian Forest Service’s Laurentian Forestry Centre for the metabarcoding and Illumina library prep.

## References

1. Neher, D. A., Weicht, T. R. & Barbercheck, M. E. Linking invertebrate communities to decomposition rate and nitrogen availability in pine forest soils. Applied Soil Ecology 54, 14–23 (2012).

2. Maab, S., Caruso, T. & Rillig, M. C. Functional role of microarthropods in soil aggregation. Pedobiologia 58, 59–63 (2015).

3. Rousseau, L. et al. Forest floor mesofauna communities respond to a gradient of biomass removal and soil disturbance in a boreal jack pine (Pinus banksiana) stand of northeastern Ontario (Canada). Forest Ecology and Management 407, 155–165 (2018).

4. Bird, G. A. & Chatarpaul, L. Effect of whole-tree and conventional forest harvest on soil microarthropods. Canadian Journal of Zoology 64, 1986–1993 (1986).

5. Battigelli, J. P., Spence, J. R., Langor, D. W. & Berch, S. M. Short-term impact of forest soil compaction and organic matter removal on soil mesofauna density and oribatid mite diversity. Canadian Journal of Forest Research 34, 1136–1149 (2004).

6. Addison, J. A. & Barber, K. N. Response of Soil Invertebrates to Clear-cutting and Partial Cutting in a Boreal Mixedwood Forest in Northern Ontario. (Canadian Forest Service, Great Lakes Forestry Centre, 1997).

7. Rousseau, L. et al. Long-term effects of biomass removal on soil mesofaunal communities in northeastern Ontario (Canada) jack pine (Pinus banksiana) stands. Forest Ecology and Management 421, 72–83 (2018).

8. Porter, T. M. & Hajibabaei, M. Scaling up: A guide to high-throughput genomic approaches for biodiversity analysis. Molecular Ecology 27, 313–338 (2018).

9. Hajibabaei, M., Spall, J. L., Shokralla, S. & van Konynenburg, S. Assessing biodiversity of a freshwater benthic macroinvertebrate community through nondestructive environmental barcoding of DNA from preservative ethanol. BMC Ecology 12, 28 (2012).

10. Yu, D. W. et al. Biodiversity soup: metabarcoding of arthropods for rapid biodiversity assessment and biomonitoring: Biodiversity soup. Methods in Ecology and Evolution 3, 613–623 (2012).

11. Oliverio, A. M., Gan, H., Wickings, K. & Fierer, N. A DNA metabarcoding approach to characterize soil arthropod communities. Soil Biology and Biochemistry 125, 37–43 (2018).

12. Watts, C. et al. DNA metabarcoding as a tool for invertebrate community monitoring: a case study comparison with conventional techniques: Monitoring invertebrates using DNA metabarcoding. Austral Entomology (2019). doi:10.1111/aen.12384

13. Hebert, P. D. N., Cywinska, A., Ball, S. L. & deWaard, J. R. Biological identifications through DNA barcodes. Proceedings of the Royal Society B: Biological Sciences 270, 313–321 (2003).

14. Hajibabaei, M., Baird, D. J., Fahner, N. A., Beiko, R. & Golding, G. B. A new way to contemplate Darwin’s tangled bank: how DNA barcodes are reconnecting biodiversity science and biomonitoring. Phil. Trans. R. Soc. B 371, 20150330 (2016).

15. Hajibabaei, M., Porter, T. M., Wright, M. & Rudar, J. COI metabarcoding primer choice affects richness and recovery of indicator taxa in freshwater systems. (Genomics, 2019). doi:10.1101/572628

16. Hajibabaei, M. et al. Watered-down biodiversity? A comparison of metabarcoding results from DNA extracted from matched water and bulk tissue biomonitoring samples. (Ecology, 2019). doi:10.1101/575928

17. Anderson, M. J. & Walsh, D. C. I. PERMANOVA, ANOSIM, and the Mantel test in the face of heterogeneous dispersions: What null hypothesis are you testing? Ecological Monographs 83, 557–574 (2013).

18. Callahan, B. J., McMurdie, P. J. & Holmes, S. P. Exact sequence variants should replace operational taxonomic units in marker-gene data analysis. The ISME Journal 11, 2639–2643 (2017).

19. Glassman, S. I. & Martiny, J. B. Ecological patterns are robust to use of exact sequence variants versus operational taxonomic units. mSphere 3, e00148–18 (2018).

20. Ficetola, G. F. et al. Replication levels, false presences and the estimation of the presence/absence from eDNA metabarcoding data. Molecular Ecology Resources 15, 543–556 (2015).

21. Porter, T. M. & Hajibabaei, M. Over 2.5 million COI sequences in GenBank and growing. PLoS ONE 13, e0200177 (2018).

22. Ratnasingham, S. & Hebert, P. D. BOLD: The Barcode of Life Data System (http://www.barcodinglife.org). Molecular ecology notes 7, 355–364 (2007).

23. Emilson, C. E. et al. DNA metabarcoding and morphological macroinvertebrate metrics reveal the same changes in boreal watersheds across an environmental gradient. Scientific Reports 7, (2017).

24. Hoage, J. F. J. Metabarcoding soil microarthropods for soil quality assessment: Importance of integrated taxonomy, phylogenetic marker selection and sampling design. (Laurentian University, 2018).

25. Kwiaton, M. et al. Island Lake Biomass Harvest Research and Demonstra on Area: Establishment Report. 82 (2014).

26. Venier, L. A. et al. Ground-dwelling arthropod response to fire and clearcutting in jack pine: implications for ecosystem management. Canadian Journal of Forest Research 47, 1614–1631 (2017).

27. Braid, M. D., Daniels, L. M. & Kitts, C. L. Removal of PCR inhibitors from soil DNA by chemical flocculation. Journal of Microbiological Methods 52, 389–393 (2003).

28. Schmidt, P.-A. et al. Illumina metabarcoding of a soil fungal community. Soil Biology and Biochemistry 65, 128–132 (2013).

29. Kennedy, K., Hall, M. W., Lynch, M. D. J., Moreno-Hagelsieb, G. & Neufeld, J. D. Evaluating Bias of Illumina-Based Bacterial 16S rRNA Gene Profiles. Applied and Environmental Microbiology 80, 5717–5722 (2014).

30. Gibson, J. et al. Simultaneous assessment of the macrobiome and microbiome in a bulk sample of tropical arthropods through DNA metasystematics. PNAS 111, 8007–8012 (2014).

31. Folmer, O., Black, M., Hoeh, W., Lutz, R. & Vrijenhoek, R. DNA primers for amplification of mitochondrial cytochrome c oxidase subunit I from diverse metazoan invertebrates. Molecular marine biology and biotechnology 3, 294–299 (1994).

32. Illumina. 16S metagenomic sequencing library preparation - Preparing 16S ribosomal RNA gene amplicons for the Illumina MiSeq System. (2013). Available at: https://support.illumina.com/downloads/16s_metagenomic_sequencing_library_preparation.html.

33. Tange, O. GNU Parallel - The Command-Line Power Tool.; login: The USENIX Magazine February, xs42–47 (2011).

34. Tedersoo, L. et al. 454 Pyrosequencing and Sanger sequencing of tropical mycorrhizal fungi provide similar results but reveal substantial methodological biases. New Phytologist 188, 291–301 (2010).

35. Kunin, V., Engelbrektson, A., Ochman, H. & Hugenholtz, P. Wrinkles in the rare biosphere: pyrosequencing errors can lead to artificial inflation of diversity estimates. Environmental Microbiology 12, 118–123 (2010).

36. R Core Team. R: A Language and Environment for Statistical Computing. (2017). Available at: https://www.R-project.org/.

37. Oksanen, J. et al. vegan: Community Ecology Package. R package version 2.5-2. (2018). Available at: https://CRAN.R-project.org/package=vegan.

38. Weiss, S. et al. Normalization and microbial differential abundance strategies depend upon data characteristics. Microbiome 5, 27 (2017).

39. Polz, M. F. & Cavanaugh, C. M. Bias in template-to-product ratios in multitemplate PCR. Applied and environmental Microbiology 64, 3724–3730 (1998).

40. Suzuki, M. T. & Giovannoni, S. J. Bias caused by template annealing in the amplification of mixtures of 16S rRNA genes by PCR. Applied and environmental microbiology 62, 625–630 (1996).

41. Elbrecht, V. & Leese, F. Can DNA-Based Ecosystem Assessments Quantify Species Abundance? Testing Primer Bias and Biomass—Sequence Relationships with an Innovative Metabarcoding Protocol. PLOS ONE 10, e0130324 (2015).

42. Ritchie, M. E. et al. limma powers differential expression analyses for RNA-sequencing and microarray studies. Nucleic Acids Research 43, e47–e47 (2015).

43. Pedersen, T. L. ggforce: Accelerating ‘ggplot2’. (2019). Available at: https://CRAN.R-project.org/package=ggforce.

44. Goral, F. & Schellenberg, J. goeveg: Functions for Community Data and Ordinations. (2018). Available at: https://CRAN.R-project.org/package=goeveg.

45. Shapiro, S. S. & Wilk, M. B. An Analysis of Variance Test for Normality (Complete Samples). Biometrika 52, 591–611 (1965).

46. Benjamini, Y. & Hochberg, Y. Controlling the false discovery rate: a practical and powerful approach to multiple testing. Journal of the Royal Statistical Society 57, 289–300 (1995).

47. Martinez Arbizu, P. pairwiseAdonis: Pairwise multilevel comparison using adonis. R package version 0.0.1. (2017). Available at: https://github.com/pmartinezarbizu/pairwiseAdonis.

48. De Cáceres, M. & Legendre, P. Associations between species and groups of sites: indices and statistical inference. Ecology 90, 3566–3574 (2009).

49. Revelle, W. psych: Procedures for Psychological, Psychometric, and Personality Research. (2018). Available at: https://CRAN.R-project.org/package=psych.

50. Wei, T. & Simko, V. R package ‘corrplot’: Visualization of a Correlation Matrix (Version 0.84). (2017). Available at: https://github.com/taiyun/corrplot.

51. Foster, Z. S. L., Sharpton, T. J. & Grünwald, N. J. Metacoder: An R package for visualization and manipulation of community taxonomic diversity data. PLOS Computational Biology 13, e1005404 (2017).

52. Chamberlain, S. & Foster, Z. taxa: Taxonomic Classes. (2018).

