## Supplementary Material for "Variations in terrestrial arthropod DNA metabarcoding methods recovers robust beta diversity but variable richness and site indicators based on exact sequence variants"

**Table S1. Read counts for all taxa**

|  | Marker |  |  |
| --- | --- | --- | --- |
| Data | BE | F230R_modN | BE+F230R_modN |
| Raw x 2 | 22,742,760 | 18,442,391 | 41,185,151 |
| Paired | 17,941,208 | 17,354,156 | 35,295,364 |
| Primer-trimmed | 16,184,320 | 17,011,213 | 33,195,533 |

**Table S2. ESV counts for all taxa**

|  | Marker |  |  |
| --- | --- | --- | --- |
| Data | BE | F230R_modN | BE+F230R_modN |
| ESVs | 50,857 | 16,769 | 67,626 |
| Reads in ESVs | 8,269,741 | 11,292,505 | 19,562,246 |
| Proportion of raw reads (%) | 36.4 | 61.2 | 47.5 |

**Table S3. Arthropoda ESV counts**

|  | Marker |  |  |
| --- | --- | --- | --- |
| Data | BE | F230R_modN | BE+F230R_modN |
| Arthropoda ESVs | 775 | 2,823 | 3,598 |
| Reads in Arthropoda ESVs | 294,070 | 2,398,638 | 2,692,708 |
| Proportion of raw reads (%) | 1.3 | 13 | 6.5 |

**Fig S1. Overview of sampling methods.** To look at the effect of increasing volume of field soil sampled, the 1C3E experiment sampled up to 36 ‘cores’ per site, each layer kept separate. To look at the effect of pooling soil samples before DNA extraction, the XC1E experiment pooled up to 15 ‘cores’ while keeping layers separate. To look at the effect of processing 1 or 3 DNA extractions per sample, the 1C1E samples were compared with the 1C3E samples that used one or three pooled DNA extractions, respectively.

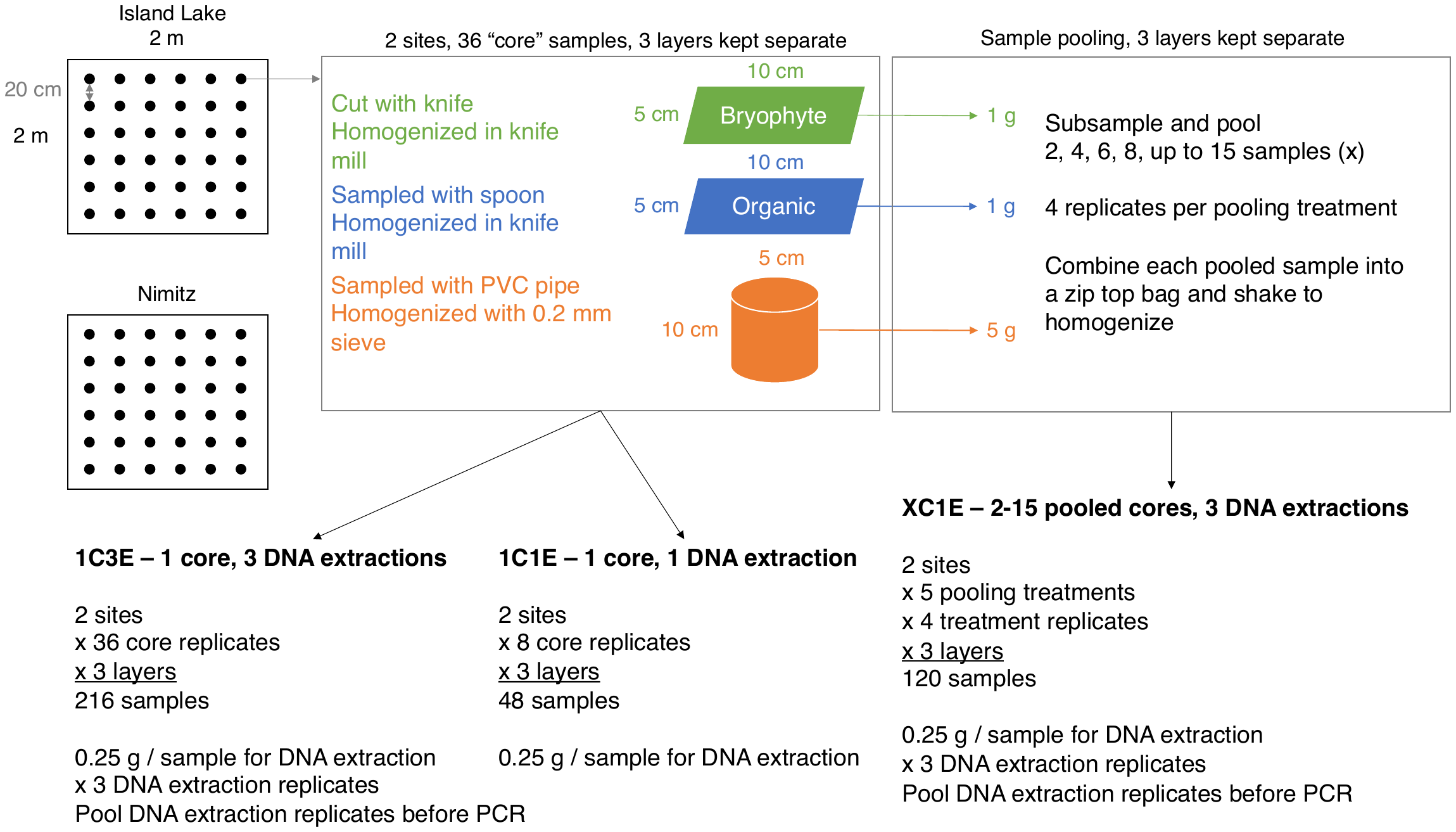

**Fig S2. Original taxonomic distribution of sequenced reads.** Results are shown by ESVs and reads. Only results from the top 10 most common phyla are shown for clarity. ESV and read numbers are labelled for only the top 3 groups for clarity.

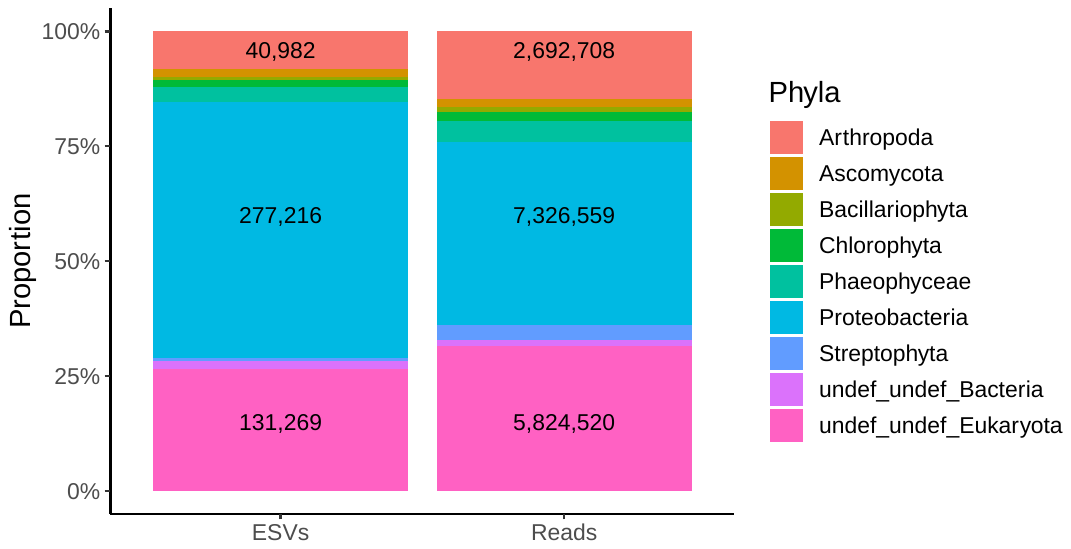

**Fig S3: A small proportion of arthropoda could be confidently identified using the current reference database.** Results are shown at the species, genus, and family ranks where minimum bootstrap support cutoffs were used to filter for confidently identified taxa.

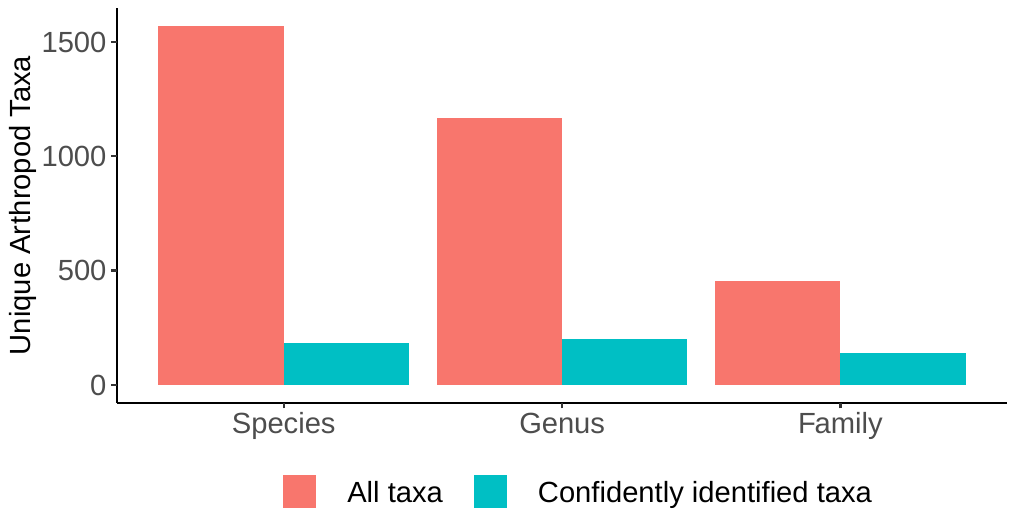

**Fig S4. Arthropoda sequencing is saturated.** Rarefaction curves from the Island lake site (ILC) (top row) and the Nimitz site (NZC) (bottom row) are shown. Reads are plotted against exact sequence variants (ESVs) for each set of experiments: 1C1E (one soil core, one DNA extraction), 1C3E (one soil core, three pooled DNA extractions), XC3E (2-15 pooled soil cores, three pooled DNA extractions). Curves are colored according to the layer they were sampled from: bryophyte layer (green), organic horizon (blue), mineral horizon (orange). The vertical guide line indicates the library size at the 15^th^ percentile.

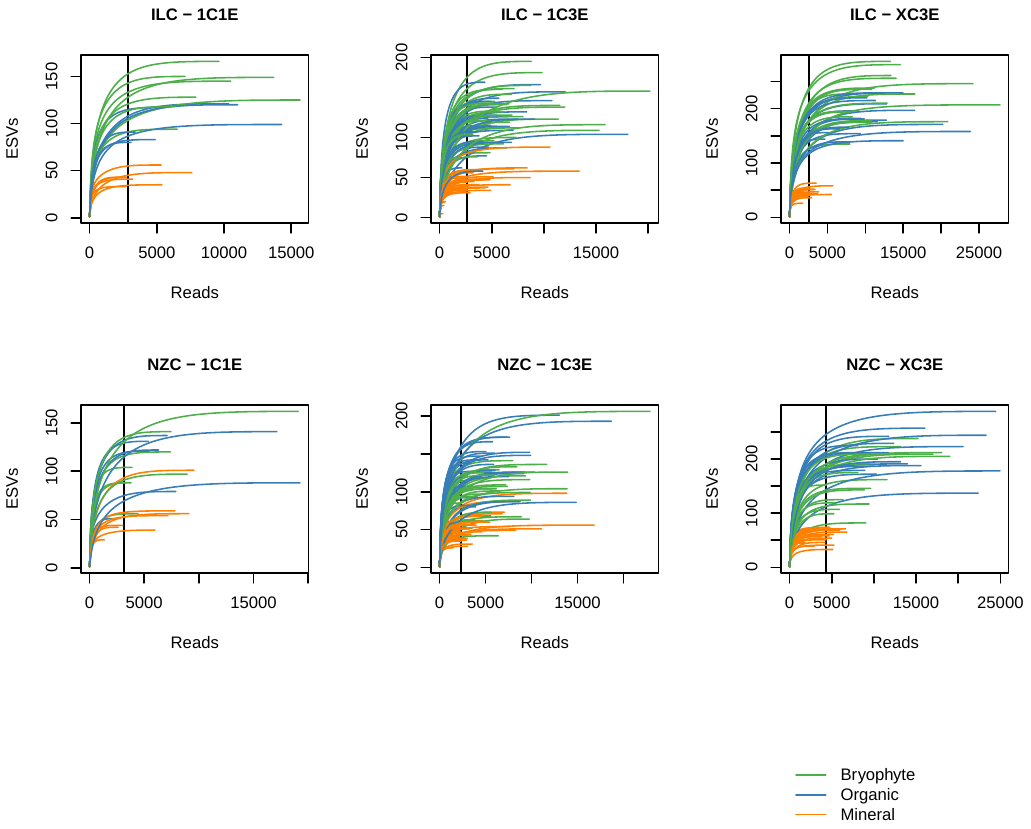

**Fig S5. Many unique ESVs are recovered from each soil layer, especially from the bryophyte and organic layers.** Based on normalized library sizes. Data pooled across all samples and methods.

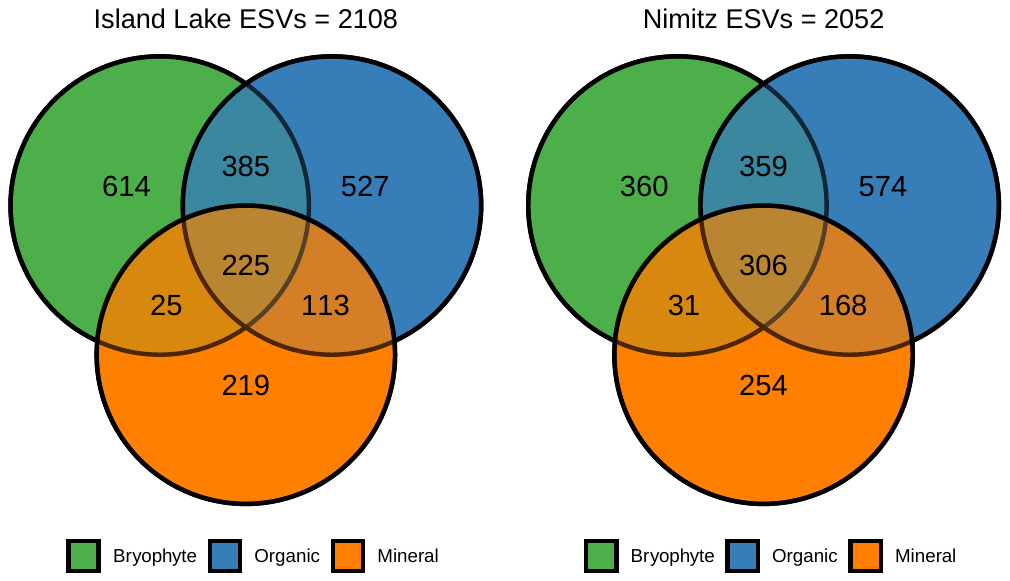

**Fig S6. Arthropod richness increases varies little when more DNA extractions are performed.** This figure is a supplement to Fig 1C in the main text. No significant differences found in the richness across sites or number of extractions. Significant differences in richness found across some soil layers (bryophyte compared with mineral, p-value = 4.1e-10; organic compared with mineral, p-value = 3.0e-10) but not when comparing the bryophyte-organic layers.

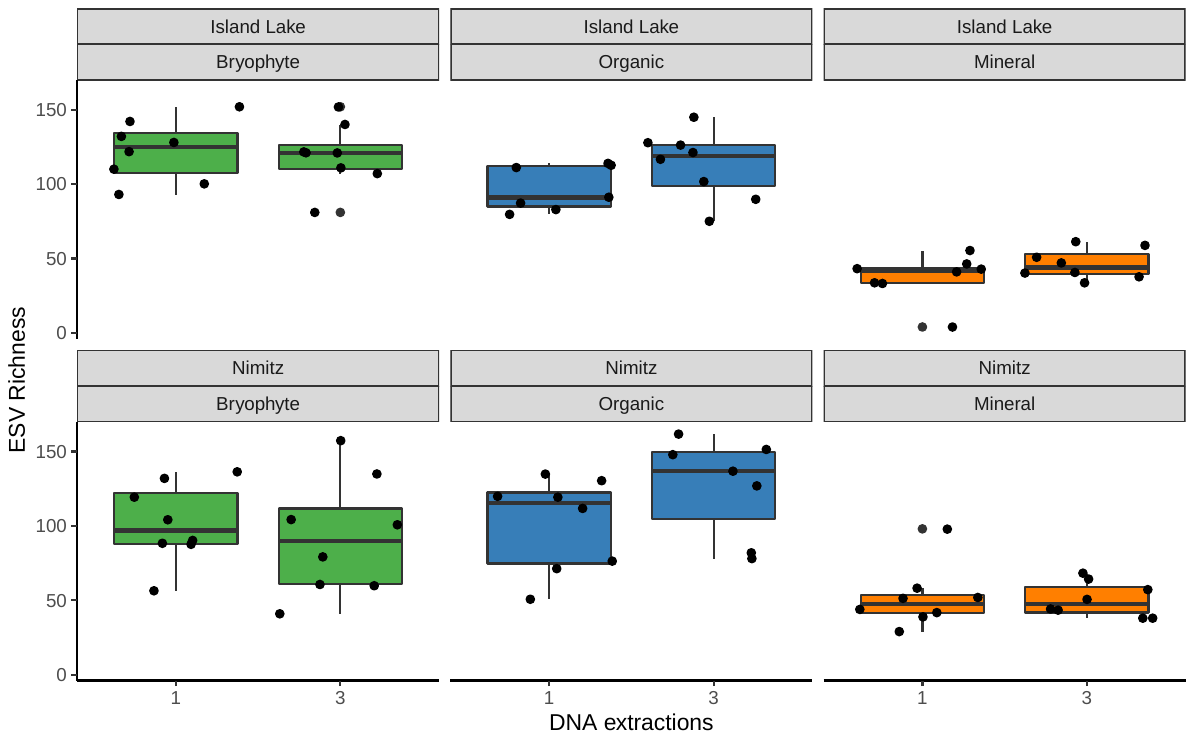

**Fig S7. Site indicators based on exact sequence variants distinguishes among two sites.** Although indicator taxa summarized to higher taxonomic ranks appear similar across sites due to our inability to confidently identify ESVs (Fig 3), sites are clearly distinguished using ESVs sampled using a variety of methods. Method abbreviations: 1C1E – one core, 1 DNA extraction; 1C3E – one core, 3 pooled DNA extractions; 2C3E – 2 pooled cores, 3 pooled DNA extractions; 4C3E – 4 pooled cores, 3 pooled DNA extractions; 6C3E – 6 pooled cores, 3 pooled DNA extractions; 8C3E – 8 pooled cores, 3 pooled DNA extractions; 915C3E – 9-15 pooled cores, 3 pooled DNA extractions. Samples from each method were subsampled down to 4 for a balanced design. For each sample, results from 3 layers (bryophyte, organic, mineral) were combined (n = 12 for each method).

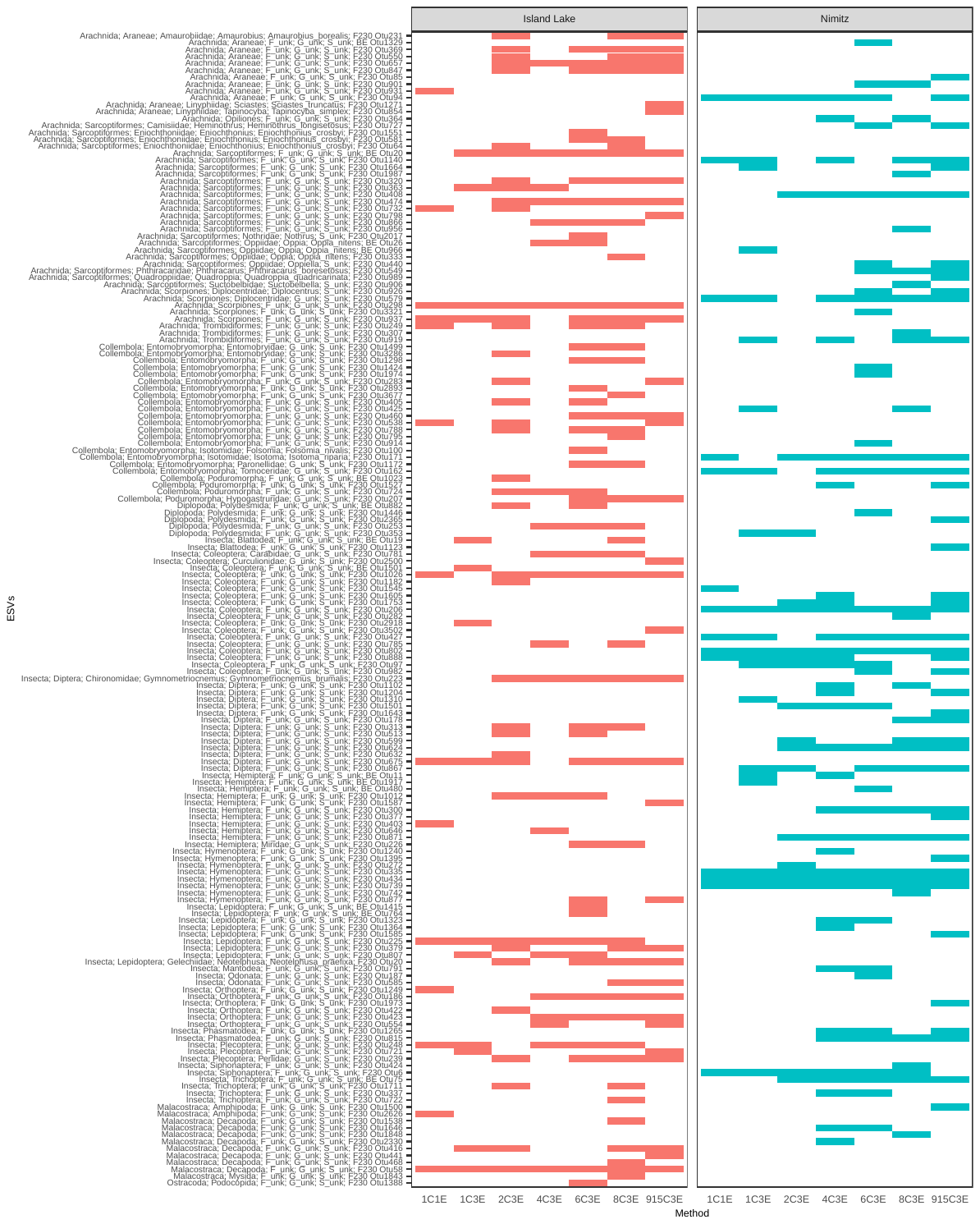

**Fig S8. Similar site indicator taxa are recovered whether 1 to 15 pooled samples are used.** Heat trees comprised of indicator taxa detected when 1 to 15 pooled samples are used. The data was pooled across both sites, then the 1C3E samples were randomly subsampled down to 4 samples per layer for a balanced comparison with the pooled core samples. In each heat tree, the frequency of indicator taxa is colored for number of pooled samples, text and node size represent ESV taxonomic assignments for the whole dataset combined. To improve readability, labels have been added only to nodes present in at least half the plotted samples. Taxa that could not be confidently identified are indicated as follows: F_unk = family unknown, G_unk = genus unknown, S_unk = species unknown.

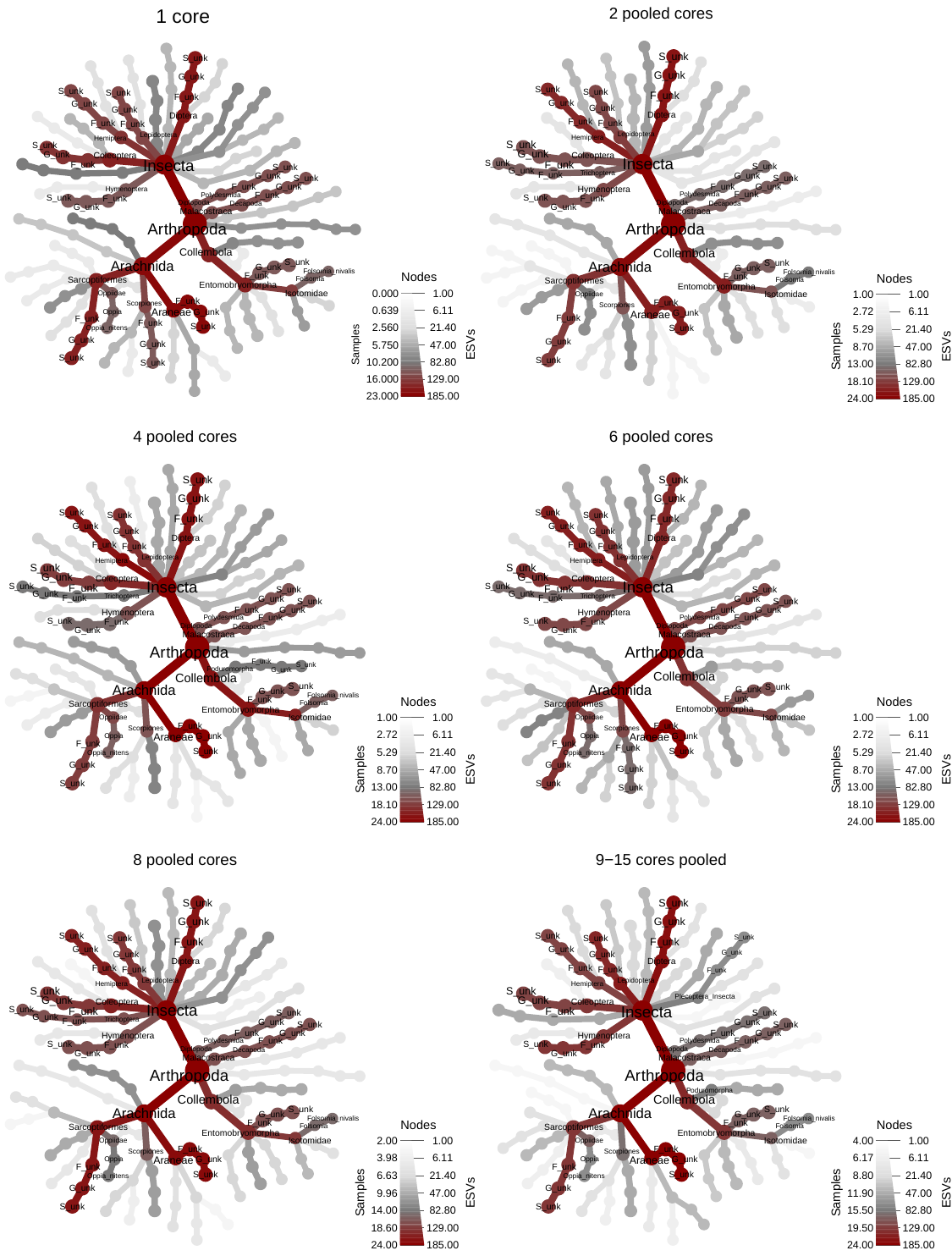

**Fig S9. Similar site indicator taxa are recovered whether 1 or 3 DNA extractions are used.** Heat trees comprised of indicator taxa detected when 1 or 3 DNA extractions are used. The data was pooled across both sites and the same 4 grid coordinates per layer were compared across the 1C1E and 1C3E methods for a balanced comparison. In each heat tree, the frequency of indicator taxa is colored for DNA extraction number, text and node size represent ESV taxonomic assignments for the whole dataset combined. To improve readability, labels have been added only to nodes present in at least half the plotted samples. Taxa that could not be confidently identified are indicated as follows: F_unk = family unknown, G_unk = genus unknown, S_unk = species unknown.

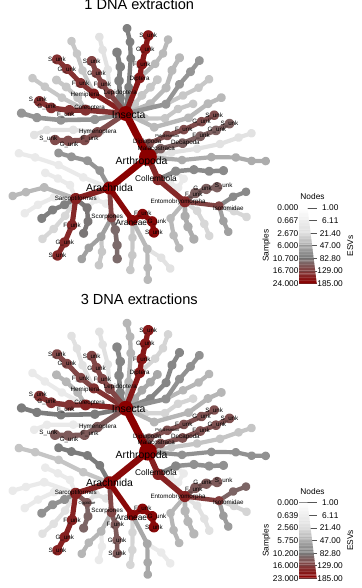

**Fig S10. Site indicator taxa are largely recovered from the bryophyte and organic layers.** Heat trees comprised of indicator taxa detected by each sampling method are shown for each site-soil stratum. For the 1C1E and 1C3E methods, samples were subsampled down to the same 4 grid coordinates for a balanced comparison across 7 methods (1C1E, 1C3E, 2C3E, 4C3E, 6C3E, 8C3E, 915C3E). In each heat tree, frequency of indicator taxa are colored for each soil stratum, text and node size represent ESV taxonomic assignments for the whole dataset combined. To improve readability, labels have been added only to nodes present in at least half the plotted samples. Taxa that could not be confidently identified are indicated as follows: F_unk = family unknown, G_unk = genus unknown, S_unk = species unknown.

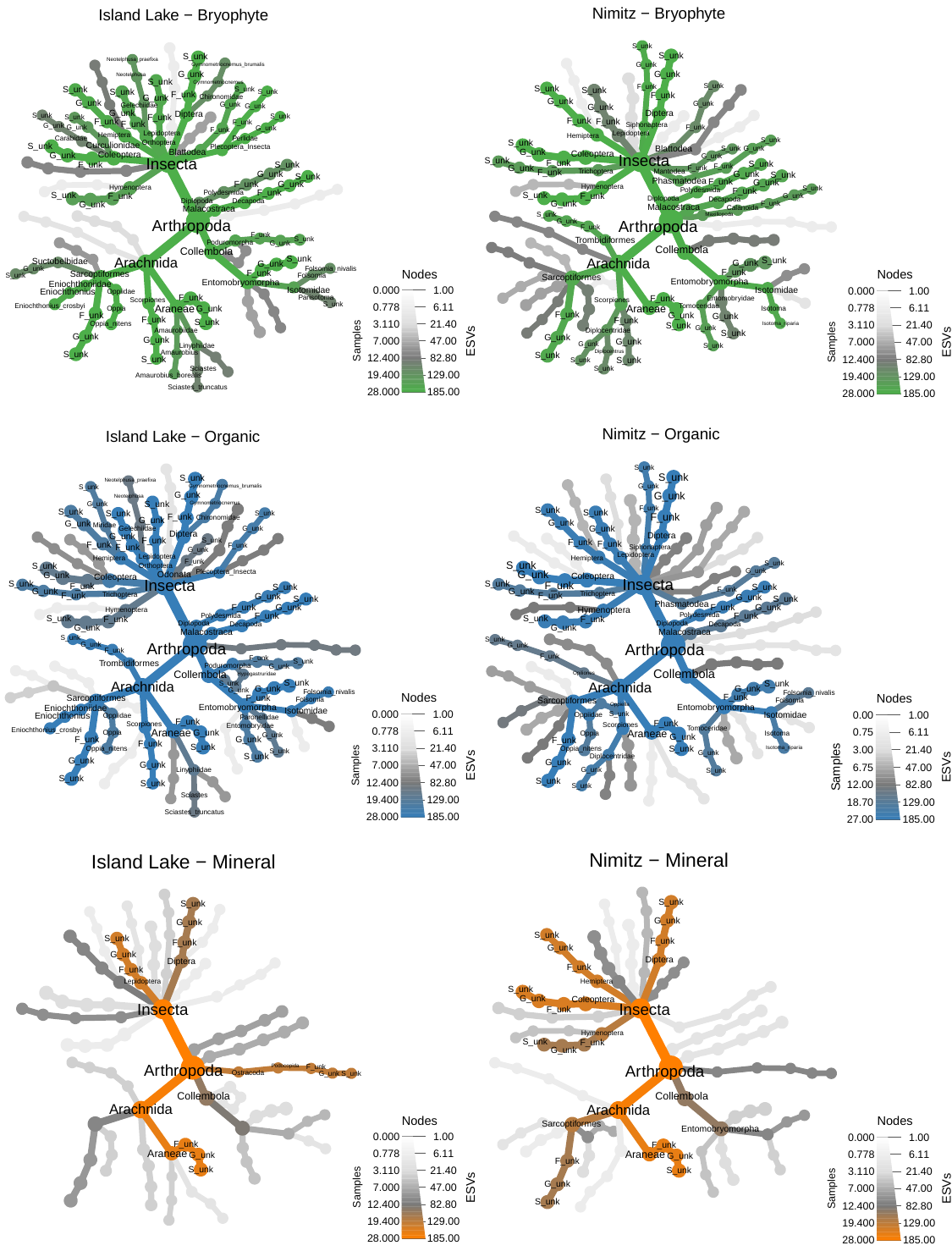
